# Predictive coding account of action perception: Evidence from effective connectivity in the Action Observation Network

**DOI:** 10.1101/722298

**Authors:** Burcu A. Urgen, Ayse P. Saygin

## Abstract

Visual perception of actions is supported by a network of brain regions in the occipito-temporal, parietal, and premotor cortex in the primate brain, known as the Action Observation Network (AON). Although there is a growing body of research that characterizes the functional properties of each node of this network, the communication and direction of information flow between the nodes is unclear. According to the predictive coding account of action perception, this network is not a purely feedforward system but has feedback connections through which prediction error signals are communicated between the regions of the AON. In the present study, we investigated the effective connectivity of the AON in an experimental setting where the human subjects’ predictions about the observed agent were violated, using fMRI and Dynamical Causal Modeling (DCM). We specifically examined the influence of the lowest and highest nodes in the AON hierarchy, pSTS and ventral premotor cortex, respectively, on the middle node, inferior parietal cortex during prediction violation. Our DCM results suggest that the influence on the inferior parietal node is through a feedback connection from ventral premotor cortex during perception of actions that violate people’s predictions.

## 1. INTRODUCTION

Over the last two decades, neurophysiological and neuroimaging studies in primates have identified a network of brain regions in occipito-temporal, parietal and premotor cortex that are associated with visual processing of actions, known as the Action Observation Network (AON, Rizzolatti & Craighero, 2004; Iacoboni & Dapretto, 2006; Saygin, 2007; Caspers et al., 2010; Nelissen et al., 2011; Saygin, 2012). Although significant progress has been made in understanding the neural correlates of action perception, one open question in the field is how the nodes of this network communicate. This question is of particular importance to be able to specify neural mechanisms that go beyond neural correlation.

There has been theoretical work that provide a neuro-mechanistic account of action perception but empirical work that directly tests it is sparse. One such model by Kilner et al. (2007a; 2007b) proposes that the AON is a predictive system, following the principles of predictive coding (Friston, 2010). In this framework, information is processed throughout the AON by means of feedforward and feedback connections, in contrast to the classic formulation of the AON, which treats action perception strictly as a feedforward process (Giese and Poggio, 2003). More specifically, the middle node of the network, parietal node has reciprocal connections between the occipito-temporal node (the lower node in the hierarchy) and the premotor node (the higher node in the hierarchy) (Figure 1), which hypothetically enables the feedforward and feedback connections within the system.

**Figure 1:**
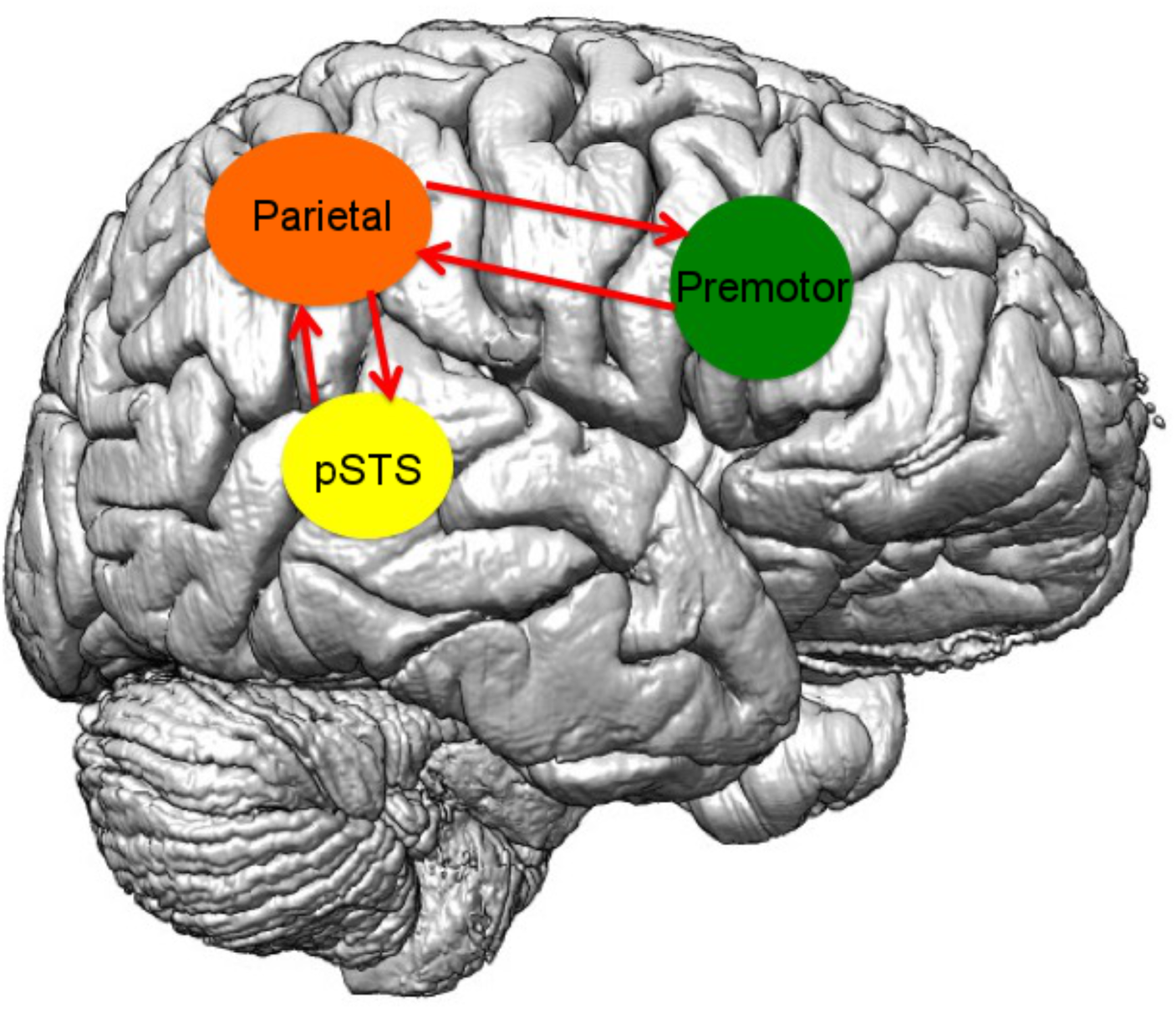
Anatomical connectivity between the core nodes of the AON.

There is indeed empirical evidence for the anatomical connectivity of the brain regions that comprise the AON. Our knowledge of the anatomical connectivity patterns in the AON comes primarily from non-human primates. In the macaque monkey, area F5 of the premotor cortex and area PF of the inferior parietal lobule have reciprocal connections (Luppino, et al., 1999). PF also has reciprocal connections with a portion of the posterior superior temporal sulcus (pSTS) that is sensitive to biological movements (Seltzer & Pandya, 1994). Analogous connectivity patterns have been proposed in the human brain (Rushworth et al., 2006; Igelstrom and Graziano, 2017).

There is also a handful of experimental studies that provide support for predictive coding account of action perception. Kilner et al. (2004), using event-related brain potentials, found that during action observation, the human brain generated a motor-preparation-like negative potential when the action was in a predictable context; no such potential was found when observation occurred within an unpredictable context. In a monkey neurophysiology study, Maranesi et al. (2014) provide direct evidence for predictive activity of visuo-motor neurons in premotor cortex and therefore it is considered to be a foundational step in supporting the predictive coding account of action understanding (Urgen and Miller, 2015). In another study, using an fMRI-adaptation paradigm, Saygin et al. (2012) found that the parietal node of the AON showed more adaptation to actions that violate predictions (via an agent who showed a mismatch between appearance and motion) than to actions that do not (via agent who shows a match between appearance and motion). The authors interpreted the differential adaptation in the parietal cortex for the prediction violations as reflecting prediction error signals generated due to a mismatch between the appearance and movement of the observed actor.

Due to the adaptation-based analysis in Saygin et al. (2012), it could not be determined whether the influence on parietal cortex came as a feedforward (bottom-up) modulation from earlier visual areas via pSTS, or as a feedback (top-down) modulation from premotor cortex in AON in the mismatch condition. The current study builds on this work and aims to reveal where that influence on parietal cortex might be generated from. Is it a top-down signal from premotor cortex or a bottom-up signal from pSTS? To address this question, we studied the effective connectivity patterns in the AON of the human brain and their modulation by the agent characteristics using functional magnetic resonance imaging (fMRI) and dynamical causal modeling (DCM) (Friston et al., 2003). Specifically, we investigated the influence of two nodes of the AON, pSTS and premotor cortex over the third node, parietal cortex and how this influence was affected by the mismatch between the appearance and motion of the observed agent.

## 2. MATERIALS AND METHODS

The fMRI data used in the present study is the same as the one collected in Urgen et al. (2019). The methodological details are provided below.

### 2.1 Participants

27 subjects (12 females, 15 males) from the student community at the University of California, San Diego participated in the study. Data of 4 subjects were not included in the analysis due to excessive head movements (3 subjects) or technical problems during data acquisition (1 subject). The subjects had no history of neurological disorders and normal or corrected-to-normal vision. Informed consent was obtained in accordance with UCSD Human Research Protections Program. The subjects were paid $25 for 1.5 hours participation in the study. All ROIs of interest for DCM analysis were identified in 18 subjects so those subjects were included in the DCM analysis.

### 2.2 Stimuli

Stimuli consisted of video clips of actions performed by 3 agents: a human agent, and the humanoid robot Repliee Q2 in two different appearances (human-like and robotic). These agents are referred here as Human, Android, Robot, respectively (Figure 2, also see Saygin et al. 2012; Urgen et al. 2013; 2019) for additional details about the stimuli).

**Figure 2.**
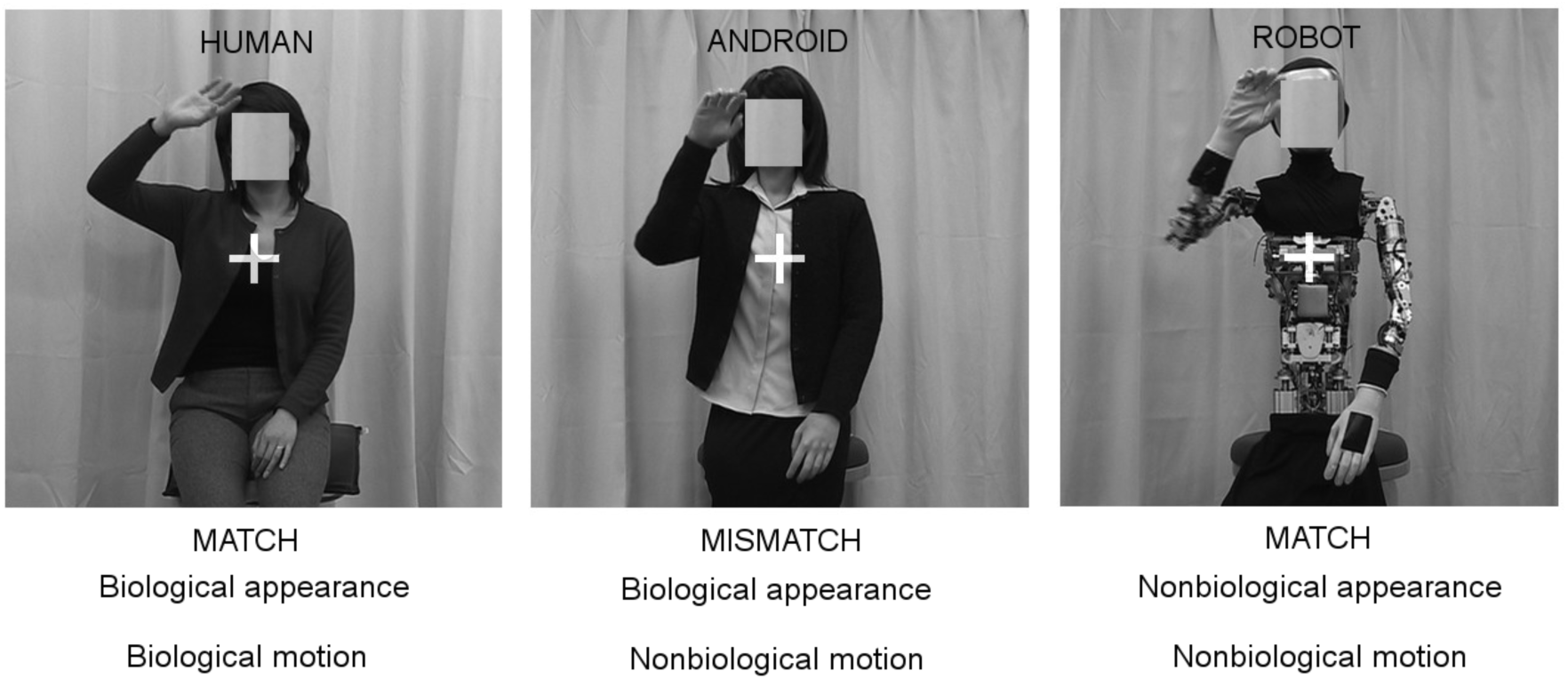
Stimuli used in the action perception experiment. There were three agents: Human with biological appearance and biological motion (match between appearance and motion), Android with biological appearance and nonbiological motion (mismatch between appearance and motion), and Robot with nonbiological appearance and nonbiological motion (match between appearance and motion).

The agents differed from each other in terms of visual appearance and motion. The Human agent had biological appearance and biological motion, the Android agent had biological appearance and nonbiological motion, and the Robot agent had nonbiological appearance and nonbiological motion. So, in this setting both Human and Robot had a *match* between their appearance and motion (both biological and nonbiological, respectively), whereas Android had a *mismatch* between the appearance and motion. All the agents performed 8 different actions. The actions were comprised of a variety of actions included drinking from a cup, grasping an object, throwing a paper, wiping a table, nudging, turning the body to the right, handwaving, and talking (for introducing herself).

### 2.3 Procedure

Each participant was given exactly the same introduction to the study and the same exposure to the videos as prior knowledge can induce biases against artificial agents (Saygin and Cicekli, 2002). Before starting fMRI scans, subjects were shown each video and were told whether each agent was a human or a robot (and thus were not uncertain about the identity of the agents during the experiment). We recorded fMRI BOLD response as subjects watched 2 sec action clips in a total of 8 runs. In each run, the experiment had a block design in which blocks consisted of video clips of one agent type (Human, Android, or Robot, see Figure 2). The experiment had 18 stimuli blocks (6 Human, 6 Android, 6 Robot) and they were presented in a pseudo-randomized order ensuring that all order combinations were presented (i.e. H-A-R, H-R-A, A-H-R, A-R-H, R-H-A, R-A-H). A rest block followed the presentation of the three blocks of agents. There, subjects fixated on a cross for a time interval varying between 8.1 sec and 13.5 sec. Each block had 9 trials (8 different actions and repetition of a randomly chosen action once) with 0.1 sec inter-stimulus interval in between the trials. Each subject was presented a different order of blocks and of stimuli within each block. In order to keep subjects’ attention throughout the experiment, a one-back task was performed in which subjects pressed a button whenever a movie was repeated.

### 2.4 Image acquisition, preprocessing and first-level analysis

We scanned our subjects at the Center for fMRI at UC San Diego using the 3T GE MR750 scanner (TR = 2.7 sec, TE = 30, Flip angle = 90, number of slices = 35, voxel size = 3mm × 3mm × 3mm, 152 volumes in each run, sequential acquisition). The stimuli were presented on a projector through a mirror mounted in the head cover in the scanner. First, the fMRI data of each subject were pre-processed with standard procedures including motion correction, slice-time correction, normalization, and smoothing using the SPM8 software. Then, two different first-level analyses were done using general linear model (GLM). In the first analysis, each agent type (Human, Android, Robot) as well as the rest blocks (fixation) were modeled as a separate condition and beta images were generated for these conditions. This analysis was done to identify the overall activity patterns and determine the ROIs of the AON. The second analysis was done to investigate the modulations in the AON via dynamical causal modeling (See Section 2.6). In the second analysis, we defined two conditions: The first condition was defined as *Actions* and consisted of actions of all three agents (Human, Android, Robot) to investigate the modulations of the connections by any action stimulus regardless of agent. The second condition was defined as the *Mismatch* condition, and consisted of the actions only by the Android, which featured a mismatch between appearance and motion, to investigate the modulations of the connections by the Mismatch condition. Motion parameters generated in the preprocessing stage were used as regressors in both analyses.

### 2.5 Identification of ROIs

We identified the ROIs of the AON by contrasting the overall activation patterns for all stimuli conditions compared to fixation (p < 0.001 uncorrected, cluster threshold k = 5 voxels) using the first first-level analysis for each subject (described in Section 2.4 above). We chose the central voxel of the activation in each node of the AON and extracted a sphere ROI that covers the activation pattern. We did this for all 18 subjects for whom we identified all ROIs of interest. The ROI time series data was then extracted using eigenvariate (threshold of p < 0.05) as a sphere with 4 mm radius.

### 2.6 Specification of the network models

In order to investigate whether a mismatch between appearance and motion of an agent during action perception was mediated through a bottom-up process from pSTS to inferior parietal cortex, or as a top-down process from ventral premotor cortex to inferior parietal cortex, we analyzed our fMRI data with dynamical causal modeling (DCM).

Dynamical causal modeling (DCM) is an effective connectivity technique to estimate the directed functional connectivity between different regions of interest with fMRI (Friston et al., 2003; Penny et al., 2004). DCM consists of two stages: Model specification and estimation, and model selection. In the model specification and estimation stage, several model architectures are specified based on the known anatomy between brain regions of interest and the hypothesis about how the connections might be modulated by the experimental manipulations. Three parameters are estimated: 1) Intrinsic connections between brain regions, 2) How the intrinsic connections are modulated by experimental manipulations, 3) The extrinsic input strength into the system. In the model selection stage, Bayesian Model Selection (BMS) procedure is used to determine the most likely model that generated the observed data. In this procedure, each model architecture in the model space is given a probability for explaining the observed data. The model that has the highest probability is then considered to be the “winning” or the most optimal model.

To test our hypothesis, we constructed 3 models that consisted of the main three ROIs of the AON, namely pSTS, the inferior parietal node, and the ventral premotor node (See the coordinates listed in Table 2) for each subject and in each hemisphere. To constrain the model space, in each of these models, the intrinsic connections between the ROIs were informed by the known anatomical connections between the regions. As such, pSTS and the parietal node, and the parietal node and the premotor node had reciprocal connections between each other (Figure 3A). Furthermore, in all models, pSTS was considered to be the node where the visual input entered the system, and all intrinsic connections were modulated by the observation of actions (*Action* condition, see Section 2.4). In other words, the observation of actions was assumed to evoke activity in pSTS first (input to the system), and then subsequently propagated to parietal and premotor cortex based on the known anatomical connections. After the first feedforward flow of information from a lower area to a higher area, feedback from a higher area to a lower area occurred in all models.

**Table 2:**
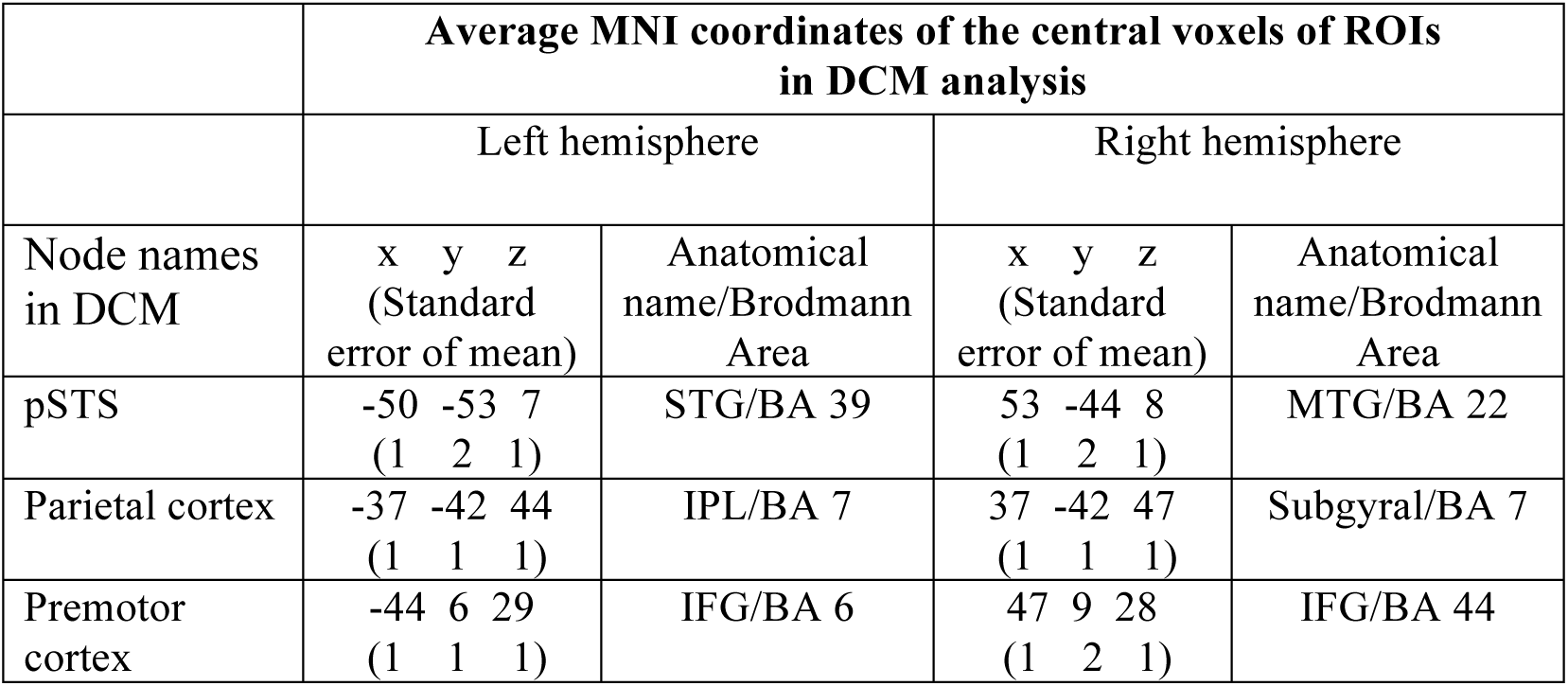
The MNI coordinates of central voxels of the ROIs used in the DCM analysis averaged across subjects. pSTS: posterior superior temporal sulcus, STG: superior temporal gyrus, MTG: middle temporal gyrus, IPL: inferior parietal lobule, IFG: inferior frontal gyrus. Values in parenthesis under x, y, z coordinates indicate the standard error of the mean.

**Figure 3:**
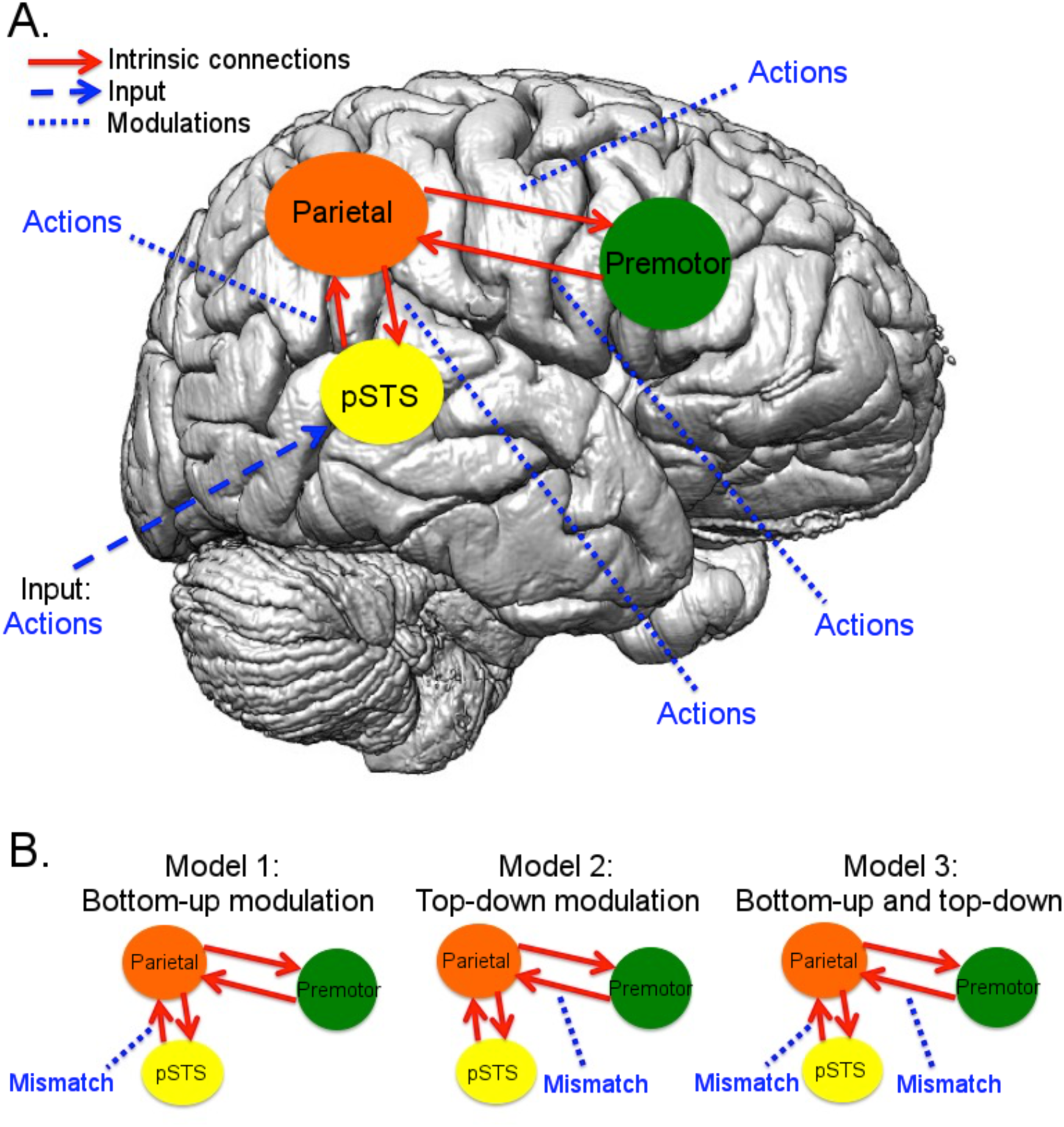
DCM models tested in the model space. (A) The DCM model that forms the basis for all tested models in the model space in (B). There are reciprocal intrinsic connections between pSTS and parietal node, and the parietal node and the premotor node informed by anatomy (red arrows). The input to the system is assumed to enter to the AON through pSTS since pSTS gets information from the visual cortex (blue dashed arrow). All the intrinsic connections are modulated by observation of actions (dashed blue lines). (B) The model space that consists of three models that correspond to our hypothesis about how the mismatch condition might modulate the intrinsic connections. Model 1 tests a bottom-up modulation from pSTS to parietal node, Model 2 tests a top-down modulation from premotor node to parietal node, and Models 3 tests both a bottom-up and a top-down modulation.

The three models differed with regard to which connections were modulated by the *Mismatch* condition (See Section 2.4, Figure 3B). The first model posits that influence on parietal cortex activity is through connections from pSTS to parietal cortex, i.e., a bottom-up modulation (Model 1). The second model posits that the influence on parietal cortex is through feedback from ventral premotor cortex, i.e., a top-down modulation (Model 2). A third possibility is that the influence would be expressed through both pSTS and premotor cortex connections (Model 3).

To identify the winning model, i.e. the model that explains that data best, Bayesian Model Selection (BMS) was used. This method determines a probability for each model, known as the exceedance probability, which is the probability that a model is more likely than any other model tested in the model space.

## 3. RESULTS

### 3.1 Brain Regions that are involved in Visual Processing of Actions

In order to identify the brain regions that were involved in visual processing of actions, we ran the contrast between the observation of actions of all agents (Human, Android, Robot) and the fixation condition. This contrast revealed the activation in early visual cortex extending dorsally to lateral occipital cortex (LOC) and ventrally to the inferior temporal cortex, as well as the core areas for AON, namely pSTS, parietal regions in the anterior part of the intra-parietal sulcus (AIP) and inferior and superior parts of the parietal lobe, and dorsal and ventral parts of the premotor cortex, all bilaterally (p < 0.001, cluster level 5) (Figure 4; Table 1 for the coordinates of all activations).

**Table 1:**
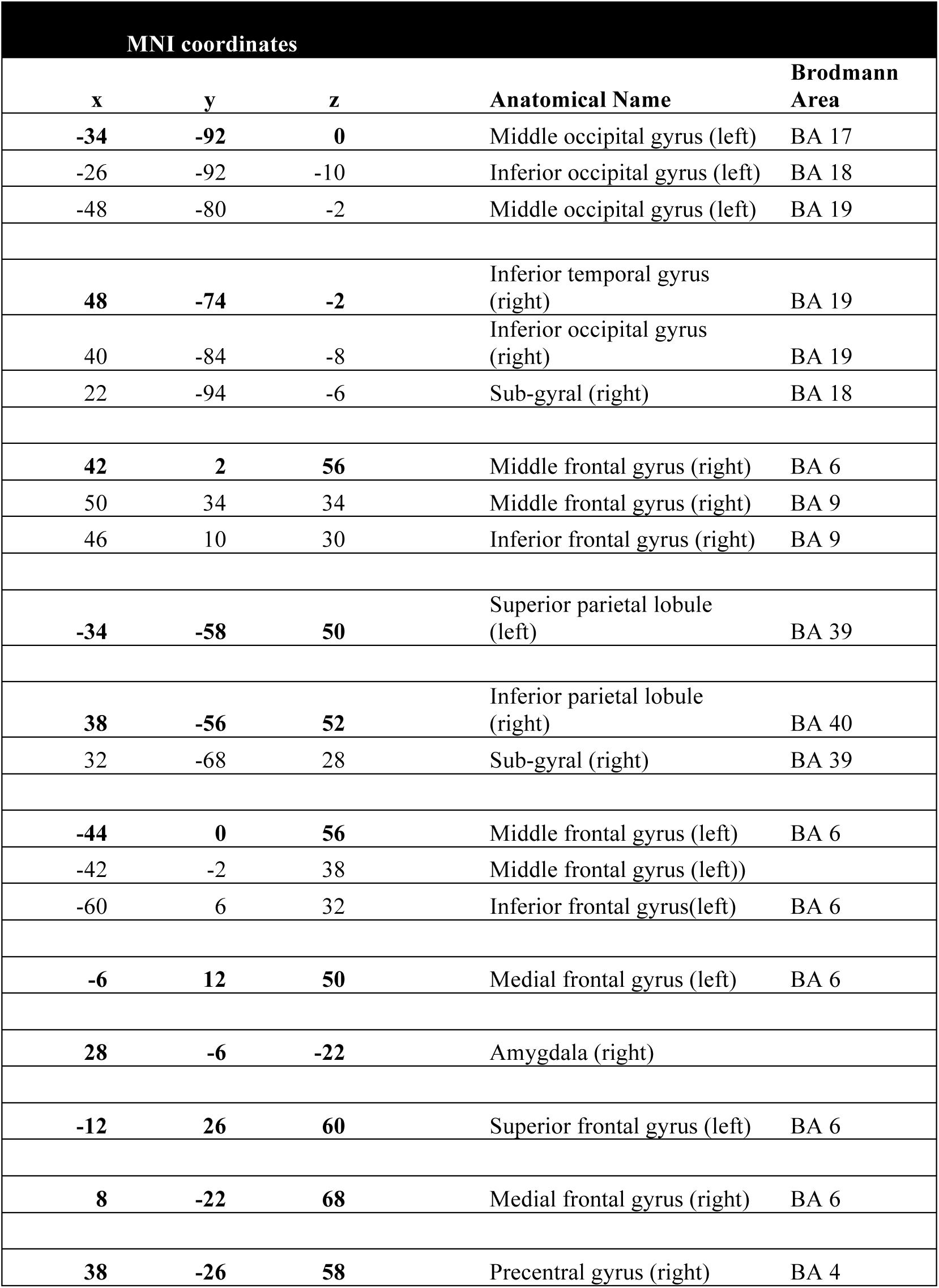
MNI Coordinates of the peak voxels of the brain regions involved in visual processing of actions based on the All Agents-Fixation contrast in the whole brain GLM analysis (Figure 3).

| MNI coordinates |  |  |  |  |
| --- | --- | --- | --- | --- |
| x | y | z | Anatomical Name | Brodmann Area |
| -34 | -92 | 0 | Middle occipital gyrus (left) | BA 17 |
| -26 | -92 | -10 | Inferior occipital gyrus (left) | BA 18 |
| -48 | -80 | -2 | Middle occipital gyrus (left) | BA 19 |
| 48 | -74 | -2 | Inferior temporal gyrus (right) | BA 19 |
| 40 | -84 | -8 | Inferior occipital gyrus (right) | BA 19 |
| 22 | -94 | -6 | Sub-gyral (right) | BA 18 |
| 42 | 2 | 56 | Middle frontal gyrus (right) | BA 6 |
| 50 | 34 | 34 | Middle frontal gyrus (right) | BA 9 |
| 46 | 10 | 30 | Inferior frontal gyrus (right) | BA 9 |
| -34 | -58 | 50 | Superior parietal lobule (left) | BA 39 |
| 38 | -56 | 52 | Inferior parietal lobule (right) | BA 40 |
| 32 | -68 | 28 | Sub-gyral (right) | BA 39 |
| -44 | 0 | 56 | Middle frontal gyrus (left) | BA 6 |
| -42 | -2 | 38 | Middle frontal gyrus (left)) |  |
| -60 | 6 | 32 | Inferior frontal gyrus(left) | BA 6 |
| -6 | 12 | 50 | Medial frontal gyrus (left) | BA 6 |
| 28 | -6 | -22 | Amygdala (right) |  |
| -12 | 26 | 60 | Superior frontal gyrus (left) | BA 6 |
| 8 | -22 | 68 | Medial frontal gyrus (right) | BA 6 |
| 38 | -26 | 58 | Precentral gyrus (right) | BA 4 |

**Figure 4:**
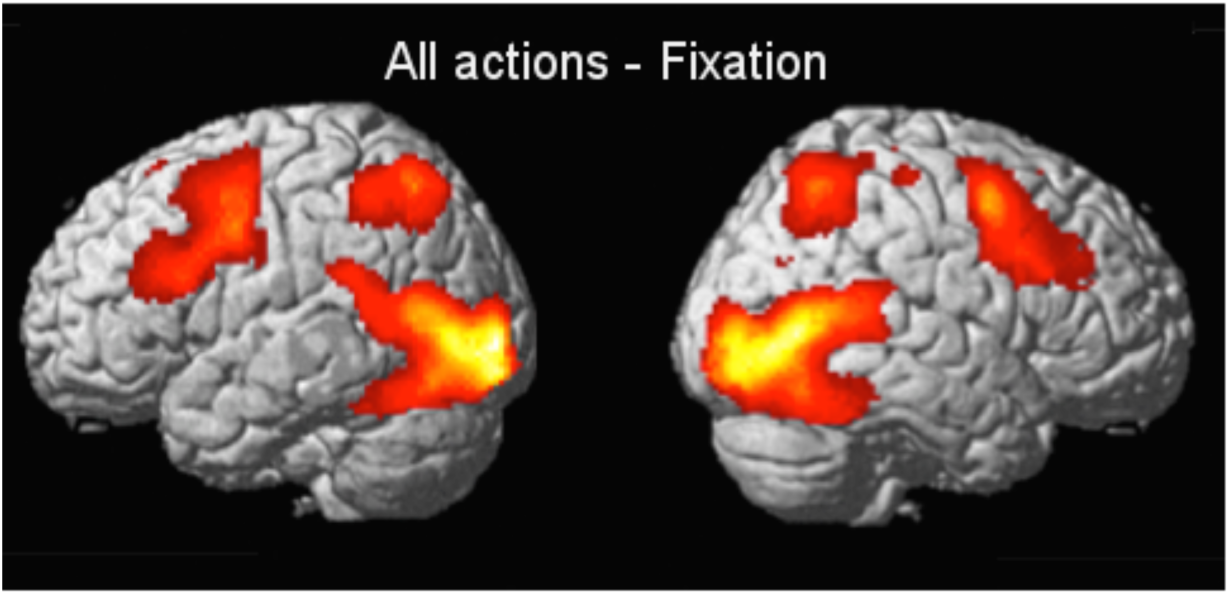
Whole brain GLM analysis with the contrast All Agents (Human, Android, Robot) – Fixation (p < 0.001, cluster threshold k = 5 voxels) at the group level. The contrast revealed activation in early visual areas extending dorsally to lateral occipital cortex (LOC), and ventrally to inferior temporal cortex, pSTS, parietal cortex, and premotor cortex dorsally and ventrally in both hemispheres. See the coordinates in Table 1.

In order to deal with the expansion of model space with increasing number of ROIs and constrain the model space used in the effective connectivity analysis, we extracted ROIs from pSTS, inferior parietal, and ventral premotor cortex in each subject, and excluded the areas in early visual areas. The coordinates of the central voxels of the ROIs averaged over subjects are displayed in Table 2. On average, subjects were 96 percent accurate in the behavioral task.

### 3.2 Effective Connectivity with DCM and Model Selection with BMS

The three DCMs included the three ROIs: pSTS, the inferior parietal node, and the premotor node (ventral premotor cortex). The intrinsic connections included the reciprocal connections between pSTS and the parietal node, and the parietal node and the ventral premotor cortex. The input into the system was considered to come from pSTS. Observation of all actions was considered to modulate all intrinsic connections (defined by the *Action* condition, see Section 2.4), and the *Mismatch* condition was considered to modulate either the pSTS-parietal connection (Model 1), premotor-parietal connection (Model 2), or both of these connections (Model 3).

BMS analysis on the three DCMs showed that Model 2 was the winning (optimal) model in both hemispheres, whose probability was 0.43 in the left and 0.50 in the right (Figure 5). The next best model was Model 1 whose probability was 0.32 on the left, and 0.36 on the right. The least likely model, Model 3 had a probability of 0.25 on the left, and 0.14 on the right.

**Figure 5:**
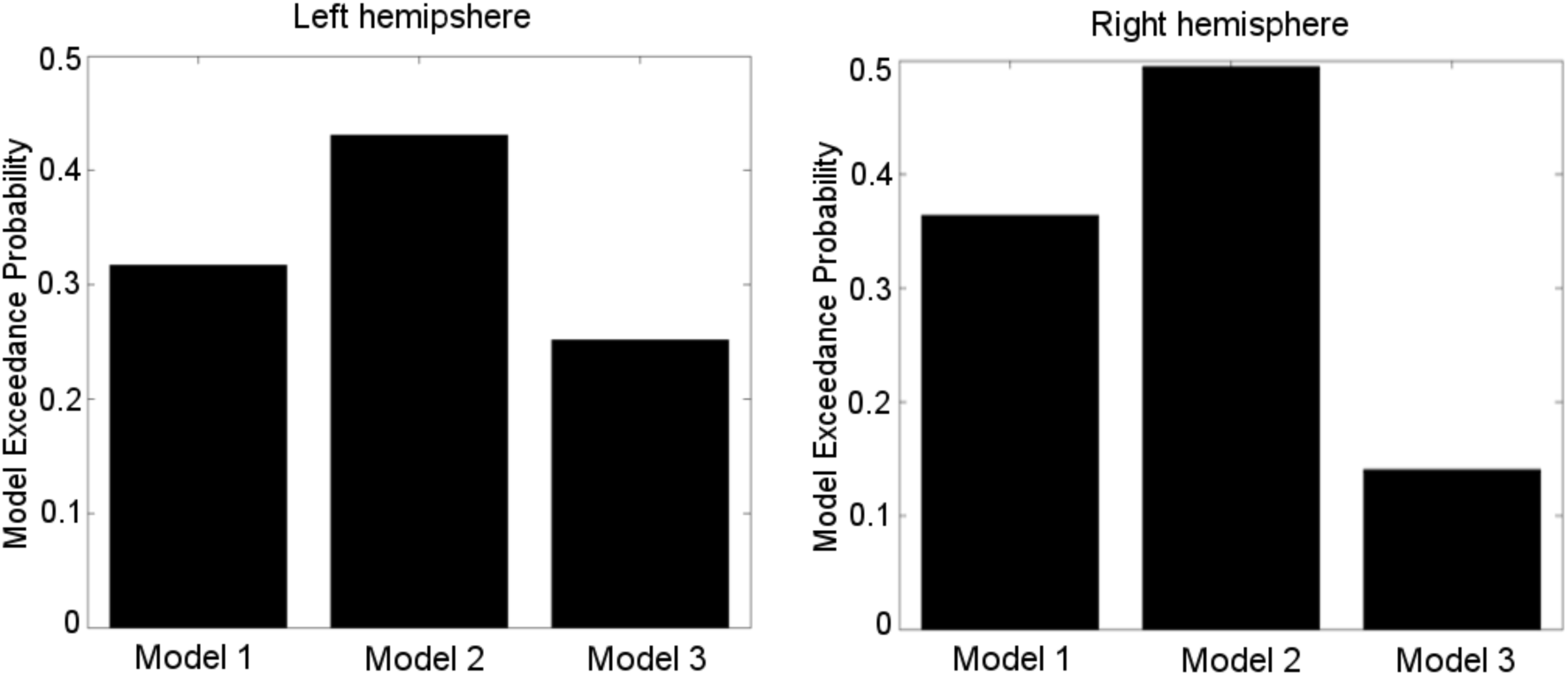
The exceedance probability of each model in the model space. Image on the left shows the results of the model testing in the left hemisphere, and the one on the right shows the results of the right hemisphere. In both hemispheres, Model 2 has the highest probability.

The intrinsic connection strengths between the ROIs in the winning model, Model 2, are listed in Table 3. All connection strengths were found to be significant (greater than 0 by a one-sample t-test, p < 0.0001) confirming the anatomical connectivity of each pair of regions. In addition, the connection directed from pSTS to the parietal node was estimated to be stronger than the other three connections (parietal to pSTS, parietal to premotor, premotor to parietal) both in the left and the right hemisphere.

**Table 3:** The parameters of the intrinsic connectivity that begins with the endogenous activity of actions in the winning model (Model 2) in both hemispheres. The values in the table indicate the mean connection strength across all subjects.

|  | FROM |  |  |  |  |  |  |
| --- | --- | --- | --- | --- | --- | --- | --- |
| TO |  | Left hemisphere |  |  | Right hemisphere |  |  |
|  |  | pSTS | Parietal | Premotor | pSTS | Parietal | Premotor |
|  | pSTS | -0.4962 | 0.0382 | - | -0.4964 | 0.0342 | - |
|  | Parietal | 0.2016 | -0.4994 | 0.0069 | 0.2168 | -0.4994 | 0.0071 |
|  | Premotor | - | 0.0449 | -0.5000 | - | 0.0461 | -0.5000 |

On the other hand, in the winning model Model 2, the modulatory effects of *Actions* on all four connections were found to be significant (greater than 0 by a one-sample t-test, p < 0.0001). The strength of the modulatory effects of *Actions* is shown in Table 4 and displayed on the right hemisphere in Figure 6 together with input strength. The connection between pSTS and the parietal node was modulated most strongly. The modulatory effect of the Mismatch condition on the premotor-parietal connection was 0.0003 on both hemispheres. The input strength was -0.0058 on the left hemisphere, and - 0.0057 on the right hemisphere.

**Table 4:** The parameters of the modulatory activity of actions in the winning model (Model 2) in both hemispheres. The values in the table indicate the mean connection strength across all subjects.

| TO | FROM |  |  |  |  |  |  |
| --- | --- | --- | --- | --- | --- | --- | --- |
|  |  | Left hemisphere |  |  | Right hemisphere |  |  |
|  |  | pSTS | Parietal | Premotor | pSTS | Parietal | Premotor |
|  | pSTS | - | 0.0095 | - | - | 0.0077 | - |
|  | Parietal | 0.0798 | - | 0.0005 | 0.0850 | - | 0.0005 |
|  | Premotor | - | 0.0121 | - | - | 0.0121 | - |

**Figure 6:**
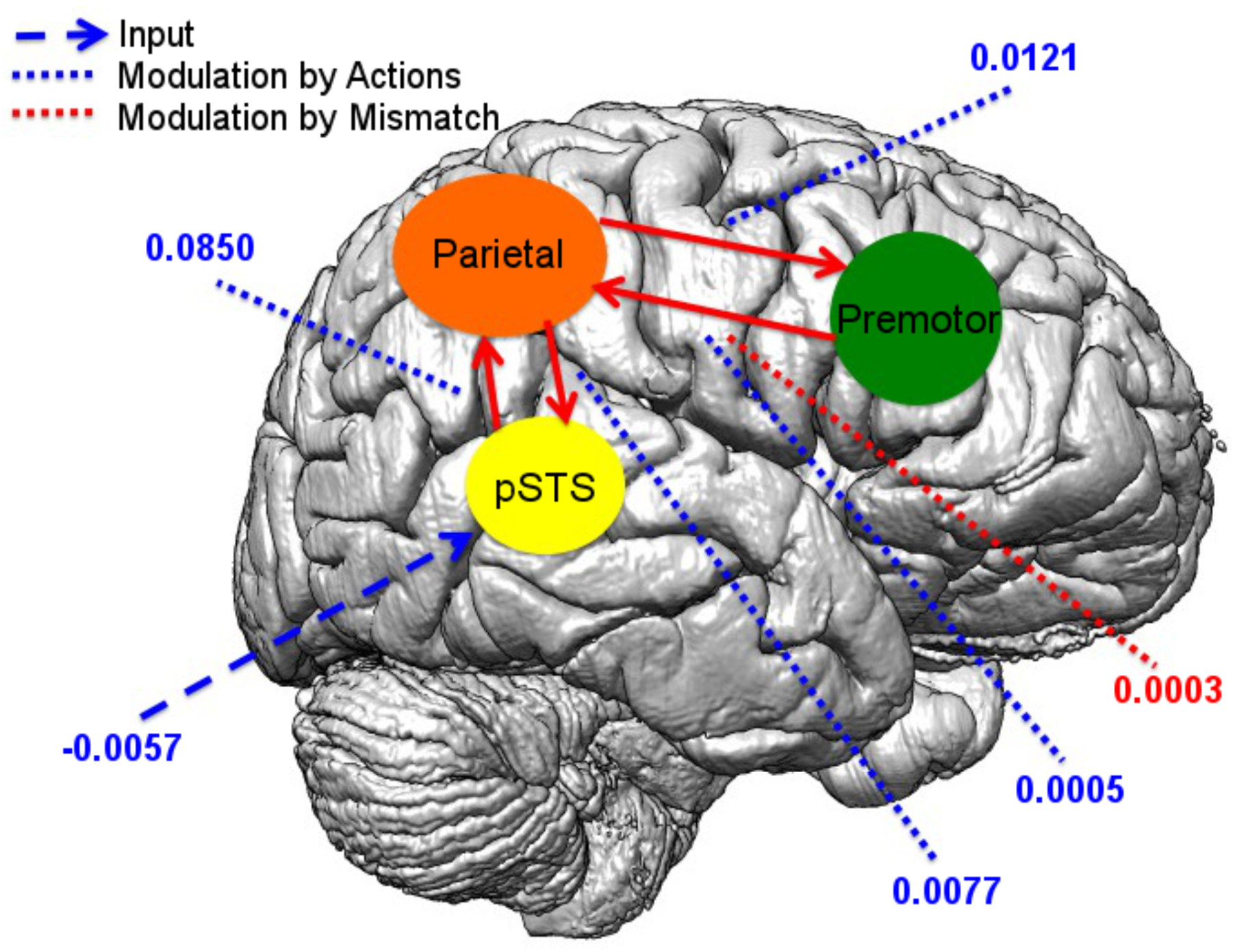
Modulatory connection strengths in the winning model, Model 2 across subjects (only right hemisphere is shown for display purposes). The mean values for action modulations for both hemispheres are also listed in Table 4. Modulations by actions are shown by the blue dashed lines. Modulation by the mismatch condition is shown by the red dashed line. All connection strengths were significantly different from 0 (with a one-sample t-test, p<0.0001).

## 4 DISCUSSION

The brain regions that are involved in visual processing of actions are relatively well-established in cognitive neuroscience (Caspers et al., 2010, Saygin, 2012). However, how the information flows between these regions have been little understood. In the present study, we aimed to estimate the effective connectivity patterns between the core nodes of the Action Observation Network and how these connections were modulated by prediction violations during action perception. Our study was primarily motivated by the findings of Saygin et al. (2012), who reported that the parietal node of the AON showed differential activity during observation of actions which were performed by an agent who possessed a mismatch between appearance and motion (a biological appearance but mechanical motion) compared to other agents who possessed a match between appearance and motion (biological appearance and motion or mechanical appearance and motion). Within the predictive coding account of action perception (Kilner et al. 2007a; 2007b), one question that has been of interest to us was whether that differential activity in parietal cortex was a top-down effect from premotor cortex or a bottom-up effect from pSTS. The current study addressed this question using fMRI and DCM.

We constructed three models to test our hypothesis. First of all, informed by well-known anatomy in the primate brain, in all these models, we created reciprocal intrinsic connections between pSTS and the inferior parietal node, and the inferior parietal node and the ventral premotor node. The input into the system was considered to enter from pSTS, which was a reasonable assumption given the involvement of pSTS in visual analysis of form and motion information in observed actions (Vangeneugden et al. 2009; 2011; Urgen et al., 2019). In addition, in all these models, we assumed that observation of actions modulated all intrinsic connections. We then constructed three specific models that corresponded to our hypotheses: A first model in which the connection from pSTS to parietal cortex was modulated, a second model in which the connection from the premotor cortex to the parietal cortex was modulated, and a third model in which both connections were modulated by the mismatch condition (the agent that exhibited a mismatch between appearance and motion). Our results show that the most likely model that best explains the data is a model in which the connection from the premotor cortex to the parietal cortex was modulated, which indicates a top-down influence.

Examination of the parameter estimates of the optimal model shows that all of the intrinsic connections were significant, confirming the well-known anatomy between these regions (Luppino, et al., 1999; Seltzer & Pandya, 1994; Rushworth et al., 2006). The strongest intrinsic connectivity was between the pSTS and the inferior parietal node. On the other hand, all of the intrinsic connections were modulated significantly by the observation of actions, which is consistent with an earlier DCM study of action observation (Sasaki et al., 2012). These results suggest that action-related information is processed via both feedforward and feedback connections in the AON. On the other hand, the mismatch between appearance and motion of an observed agent during action perception was likely mediated primarily via a feedback connection from the premotor to the parietal node of the AON.

These results provide support for the predictive coding account of action perception (Kilner et al., 2007a; 2007b), which proposes that there are reciprocal connections between the three levels of the AON, and prediction error signals are communicated via the feedback connections. Our data shows that actions are not only processed by a network that has feedforward and feedback connections consistent with previous work (Sasaki et al., 2012; Maffei et al. 2015; Gardner et al., 2015; Cardellicchio et al., 2018; Sokolov et al., 2018) but also provides evidence that in the case of prediction violations (e.g. when a moving agent exhibits a mismatch between appearance and motion), the premotor node of the AON likely sends a feedback signal to the inferior parietal node in the lower part of the hierarchy, which might indicate a prediction error signal (Saygin et al., 2012).

However, some caveats must be noted. BMS is a Bayesian approach, which assigns a probability to each model tested in the model space and constrained by the experimental design. Future studies should test the generality of this model during action observation under different task demands. For instance, the mismatch between the two visual cues, appearance and motion is a particular case where predictions are violated during action perception. Novel prediction paradigms in which there are mismatches between different visual cues, multi-sensory cues, or even cognitive cues (e.g. as in Costantini et al., 2005; Koelewijn et al., 2008; Stapel et al., 2010) should be tested in future studies of action observation with effective connectivity techniques.

## Acknowledgments

This study was supported by Qualcomm Institute, Kavli Institute for Brain and Mind, and DARPA. The authors also would like to thank Edward Nguyen for his help in fMRI data collection.

